# Increased Aperiodic Exponents Track Depression Symptom Severity

**DOI:** 10.64898/2025.12.17.694966

**Authors:** Mark R. Libowitz, Wendy Sun, Rikki Rabinovich, Jingnan Du, Justin M. Campbell, Rhiannon L. Cowan, Niloufar Shahdoust, T. Alexander Price, Tyler S. Davis, Randy L. Buckner, Shervin Rahimpour, Brian J. Mickey, Elliot H. Smith, Ben Shofty

## Abstract

Roughly one-third of patients with major depressive disorder (MDD) fail to respond to standard treatments and develop treatment-resistant MDD. For these patients, alternative therapies offer additional options but yield inconsistent outcomes. Progress has been limited by the absence of objective, brain-based biomarkers to guide target selection or track therapeutic response in real time. Instead, clinicians rely on behavioral assessments that evolve slowly over weeks to months, obscuring the underlying neural dynamics of symptom changes. Here, we test whether the aperiodic exponent of intracranial EEG (iEEG) local field potentials can serve as a neurophysiological marker of depression symptom severity.

We leveraged a large iEEG cohort (N = 20) undergoing invasive monitoring for refractory epilepsy, yielding over 1,800 contacts spanning cortical and subcortical zones. For each contact, we estimated the aperiodic exponent (thought to reflect aspects of cortical excitability) of the power spectrum across 10-100 Hz within local brain regions and across distributed cortical association networks. Depressive symptoms were assessed with the Beck Depression Inventory-II (BDI-II) immediately before intracranial resting state recordings. With respect to the BDI-II scale, participants were identified as experiencing minimal (BDI-II ≤ 13) or elevated depression symptoms (BDI-II ≥ 14). Associations between symptom severity (BDI-II total score and Somatic-Affective, Cognitive, and Anhedonia subscales) and region- or network-level exponents were modeled with ordinary least squares (OLS) regression.

The whole-brain, mean aperiodic exponent for each participant discriminated symptom status (AUC = 0.82). At the regional level, the orbitofrontal cortex, anterior cingulate cortex, insula, and amygdala showed higher exponents in the elevated depression symptom group (d = 1.18-1.71; p = 0.032-0.004). A post-hoc classification analysis across these four regions misclassified one participant per group (AUC = 0.86; 95% CI 0.64-1.00). In continuous analyses, BDI-II scores correlated positively with exponents in these same four regions (pFDR = 0.019-0.027; partial r=0.61–0.70) and at the network level in the Salience network (pFDR = 0.024; partial r = 0.63) and Default (pFDR = 0.046; partial r = 0.55) network. The Salience network significantly tracked Anhedonia symptoms (p = 0.004; partial r = 0.62).

Here we report that intracranial aperiodic exponents within fronto-limbic and insular circuits, overlapping with networks implicated in contemporary accounts of depression pathophysiology, differentiate depressive symptom status and scale with severity. These findings support the aperiodic exponent as a candidate neurophysiological marker of current depression symptom burden, with potential relevance for individualized neuromodulation in MDD.

## Introduction

Major depressive disorder (MDD) affects more than 8% of adults in the United States^1^ and is associated with increased mortality and disability compared to the general population.^2, 3^ Despite available treatments, only about one-third to one-half of patients remit with an initial antidepressant trial,^4, 5^ and approximately one-third ultimately meet criteria for treatment-resistant MDD after failing multiple interventions.^5, 6^

For these patients, alternative pharmacological or neuromodulation therapies are available, but clinical benefit remains highly variable across trials and individuals.^7–10^ Key barriers to developing more effective therapeutics are heterogeneous anatomical targeting and the absence of validated brain-based biomarkers. This motivates the search for a scalable, interpretable biological readout of current depression symptom severity that can be tracked during treatment, rather than relying solely on behavioral shifts that often accrue over weeks to months.^11, 12^ Such a marker could be used to guide treatment adaptation and target selection, ultimately improving outcomes across clinical settings.

While prior work has often emphasized the oscillatory components of neural power spectra as correlates of cognitive function and dysfunction, accumulating evidence suggests that non-oscillatory features carry physiological relevance in neurological and psychiatric conditions (e.g., Alzheimer’s, ADHD, autism, schizophrenia, and depression).^13^ Thus, in this study, we examine the aperiodic exponent of the neural power spectrum as a candidate biomarker of depression symptom severity. The aperiodic component is the scale-free, non-oscillatory background of the neural power spectrum that follows a 1/f^χ^ decay (f = frequency, χ = aperiodic exponent). In this framework, larger χ values correspond to steeper spectral slopes. Converging evidence suggests that the exponent reflects aspects of cortical excitability—potentially indexing the balance between synaptic excitation and inhibition, as well as synaptic integration timescales and membrane filtering properties—with steeper slopes generally associated with relatively greater inhibitory tone.^14, 15^ Therefore, as a single, interpretable measure of low-to-high frequency power balance that may approximate global excitability, the aperiodic exponent represents a compelling emerging candidate biomarker for real-time, individualized monitoring of brain states during treatment.

To date, three intracranial electrophysiology studies (DBS local field potentials and/or stereo-EEG) have examined the aperiodic exponent in relation to depression in distinct, clinically implanted cohorts with treatment-resistant depressive illness. Two reported that larger exponents accompanied greater severity,^16, 17^ whereas one found that larger exponents after treatment were linked to symptom improvement.^18^ These reports underscore the aperiodic exponent’s potential as a biological index of depression symptom severity but were limited by small sample sizes (N = 4–6 patients per study) and anatomically constrained sampling—either restricted to a single DBS target (subcallosal cingulate cortex or habenula) or confined to a narrower set of intracranial recording sites. Because mood fluctuations may be predicted by distributed network activity spanning multiple frequency bands, inter-regional and across-frequency interactions merit consideration beyond isolated spectral changes.^19, 20^ Of particular interest for depression is the Salience network (SAL), which was shown to be topographically expanded in patients with depression, even predicting future symptoms of anhedonia.^21^ Furthermore, SAL likely overlaps with modern neuromodulation targets, further emphasizing SAL’s role in depression.^21–23^ In addition to SAL, other networks have also been implicated in the pathophysiology of depression, including approximations of the spatially adjacent cingulo-opercular network^24^ (CG-OP), the default network^25^ (DN), and the frontoparietal control network^26^ (FPN).

In this study, we leveraged a cohort of epilepsy patients implanted with intracranial EEG (iEEG) for epilepsy monitoring (N=20), affording rare, high-precision recordings from intracranial electrodes distributed across cortical and subcortical zones. Given the frequent occurrence of depressive symptoms in this population^27^ and broad intracranial electrode coverage across frontal, limbic, insular, and temporal regions spanning association networks implicated in depression, we tested whether the aperiodic exponent covaries with concurrently measured depression symptom severity. To assess anatomical specificity, we focused on the nine regions that had consistent coverage across patients: orbitofrontal cortex (OFC), anterior cingulate cortex (ACC), dorsolateral prefrontal cortex (dlPFC), insula, amygdala, hippocampus, middle temporal gyrus (MTG), superior temporal gyrus (STG), inferior temporal gyrus (ITG). To assess distributed network-level effects, we focused on six cortical association networks: DN-A, DN-B, FPN-A, FPN-B, SAL, CG-OP.^28^ We hypothesized that larger aperiodic exponents (steeper 1/f^χ^ slopes) would track greater depression symptom severity, most prominently within corticolimbic and insular circuits.

## Materials and Methods

### Study Overview

Patients undergoing iEEG for clinical monitoring (N=20) were recruited for this study. After electrode implantation, participants completed the Beck Depression Inventory-II (BDI-II; a 21-item self-report measure of depression severity)^29^ immediately before a five-minute resting-state local field potential (LFP) recording. We then calculated the aperiodic exponent across distributed regions and association networks to evaluate its relationship with depression symptom severity.

### Participants

Twenty adult patients underwent neurosurgery at the University of Utah in Salt Lake City, UT and received intracranial monitoring with iEEG to localize seizure foci at the Epilepsy Long-Term Monitoring Unit (Table 1). This study was approved by the Institutional Review Board of the University of Utah (IRB #0011423), and all patients signed an informed consent form prior to participation. Participant eligibility included age ≥18, ability to provide informed consent, English as first language, IQ ≥ 70, no history of a resective surgery, and no significant medical comorbidities. No exclusion was made concerning sex, gender, race, or ethnicity.

**Table 1.** Participant demographics and clinical characteristics. Electrode counts reflect contacts retained after quality control. Depressive symptoms were assessed with the BDI-II scale. Seizure onset zone (SOZ) was determined by an attending neurologist following intracranial epilepsy monitoring. Abbreviations: M/F = male/female; R/L = right/left, *MTL = medial temporal lobe, PHC = parahippocampal cortex, Hipp = hippocampus, TT = temporal tip, Amy = amygdala, STG = superior temporal gyrus, MTG = middle temporal gyrus.

| ID | Age | Sex | Electrode Count | BDI-II Score | Seizure Onset Zone |
| --- | --- | --- | --- | --- | --- |
| P01 | 49 | M | 70 | 2 | R MTL |
| P02 | 27 | F | 101 | 12 | R Parietal-Occipital Cortex |
| P03 | 29 | M | 71 | 9 | R MTL |
| P04 | 22 | F | 106 | 44 | Bilateral PHC and Hipp |
| P05 | 48 | F | 70 | 28 | L Hipp |
| P06 | 26 | M | 113 | 5 | R Parietal, Occipital, Precuneus |
| P07 | 40 | F | 102 | 4 | R Hipp |
| P08 | 27 | M | 88 | 42 | Bilateral MTL |
| P09 | 46 | F | 73 | 27 | L MTL and L TT |
| P10 | 23 | F | 92 | 12 | R MTL |
| P11 | 67 | M | 81 | 17 | R Hipp |
| P12 | 24 | F | 71 | 35 | L Hipp |
| P13 | 35 | M | 101 | 9 | L Amy, R Hipp |
| P14 | 37 | F | 86 | 21 | L STG, L MTG, L TT |
| P15 | 36 | M | 96 | 19 | L MTL |
| P16 | 25 | M | 100 | 8 | Bilateral MTL |
| P17 | 18 | M | 67 | 25 | Bilateral MTL |
| P18 | 35 | M | 89 | 2 | Bilateral MTL |
| P19 | 37 | M | 140 | 3 | Bilateral MTL |
| P20 | 36 | F | 87 | 12 | Bilateral MTL |

### Intracranial Electrode Implantation and Recording

Decisions regarding electrode implantation sites were made exclusively on clinical grounds during a multidisciplinary case conference and were not influenced by, or tailored to, the present research protocol. Each patient was implanted with clinical macroelectrodes (mean ± S.D = 90.20 ± 15.15 contacts per participant) to record LFP using a 128-channel neural signal processor (Blackrock Microsystems, Salt Lake City, UT, USA), sampled at 1kHz. Electrode contacts were referenced to an intracranial contact located within the white matter.^30^

### Experimental Design

Immediately before intracranial recording sessions, participants completed the BDI-II and then underwent a five-minute resting-state recording while fixating on a gray screen with a white central crosshair. The BDI-II was used to classify participants as experiencing minimal depression symptoms (BDI-II ≤ 13) or elevated depression symptoms (BDI-II ≥ 14; mild to severe depression symptoms). Beyond the total BDI-II score, we analyzed three continuous symptom dimensions: Cognitive, Somatic–Affective, and Anhedonia subscales as validated in prior work.^31–33^

### Post-Implantation Electrode Localization, Registration, and Anatomical Labeling

Utilizing custom MATLAB software and SPM12,^34^ each patient’s preoperative T1 MRI and post-operative CT were co-registered. To localize electrodes, we leveraged the open-access Localize Electrodes Graphical User Interface (LeGUI) software, which performs co-registration of pre-operative MRI and post-operative CT sequences, image processing, normalization to Montreal Neurological Institute (MNI) brain atlas space, and automated electrode detection.^35^ Brain regions were identified using the Brainnetome atlas,^36^ grouped into regions of interest, and confirmed by two attending neurosurgeons (BS and SR). To ensure adequate coverage across subjects, we excluded regions sampled in fewer than 60% of participants. The following regions were carried forward for regional analyses: OFC, ACC, dlPFC, insula, amygdala, hippocampus, MTG, STG, and ITG.

Electrode contact coordinates in MNI space were also assigned a functional network label, based on the DU15NET-Consensus atlas.^28^ This atlas is a 15-network parcellation derived from intensively sampled individuals and includes a separate label for uncertain / low-confidence zones, which is an explicit concession to limitations of fMRI signal in certain regions. Each contact’s MNI coordinates were localized to the closest voxel in the MNI152 1mm brain volume. Then, a neighborhood of all voxels within a 3mm radius was sampled, and the most common (modal) network label within this neighborhood was identified (see Supplemental Materials for additional details). Network analyses focused on six distributed association networks based on previous work investigating these networks role in depression: DN-A, DN-B, FPN-A, FPN-B, SAL, CG-OP.^21–23^ All networks of interest had sampling in greater than 60% of participants. Electrode locations were visualized in MNI-space using the Visbrain software package.^37^

### Intracranial Electrophysiological Signal Processing

Preprocessing was performed in MNE-Python.^38^ Contacts in patient seizure onset zones, as determined by an attending clinical neurologist, were removed from the analysis, and all intracranial recordings were visually inspected to identify noisy or dead channels, which were excluded. All exclusions were settled upon prior to additional hypothesis-directed analysis. High-pass (0.5 Hz) and low-pass (100 Hz) finite impulse response filters were then applied, followed by a common average re-referencing and downsampling to 500 Hz.

Channel-wise power spectral density was estimated using Welch’s method with 2-second windows (50% overlap; zero-padded to an effective window length of 4.096 s). This produced one PSD estimate per contact across the full 5-minute recording. Spectra were resampled to a mixed log-linear frequency grid (higher relative resolution below 25 Hz), and 60-Hz line noise was attenuated by removing 58-62 Hz and replacing those values via interpolation from adjacent frequency bins. The cleaned power spectra were subsequently re-interpolated onto a linearly spaced frequency axis of 200 points spanning 10-100 Hz for spectral parameterization.

### Aperiodic Exponent Calculation

To calculate the aperiodic exponent for each contact, the fitting oscillations & one over f (FOOOF) algorithm (version 1.1)^39^ was used to parameterize neural power spectra (**Fig. 1**). We focused our analysis on the 10-100 Hz range, a frequency band well captured by iEEG and suited for broad-band analyses.^40^ Model parameters were optimized per channel via a grid search over peak width limits (minimum width set to twice the mean frequency resolution of the 10–100 Hz spectrum, 0.905 Hz; maximum widths of 2, 4, 6, 8, or 12 Hz), peak height (0.1, 0.2, 0.3), number of peaks (2, 4, 6, or 8), and aperiodic mode (“fixed” or “knee”) to minimize the mean absolute error (MAE) fit metric. All model parameters were finalized prior to data analysis. Fits with an MAE greater than 0.05 were excluded as poor model fits; MAE values for retained spectra were low and tightly distributed (mean/median = 0.023/0.022). Exponent estimates outside the range of 1.5–4.5 were removed as physiologically implausible for iEEG, consistent with the literature, which reports exponents typically ranging from 2 to 4 when fitted across 10–100 Hz.^41, 42^ Additionally, outliers in aperiodic exponent estimates were removed within each region using an interquartile range (IQR)-based criterion: values greater than 3.0 x IQR beyond the 25^th^ or 75^th^ percentile were excluded. This conservative procedure was applied per region and network to account for variability in exponent distributions (in total, only two ACC contacts were removed).

**Figure 1.**
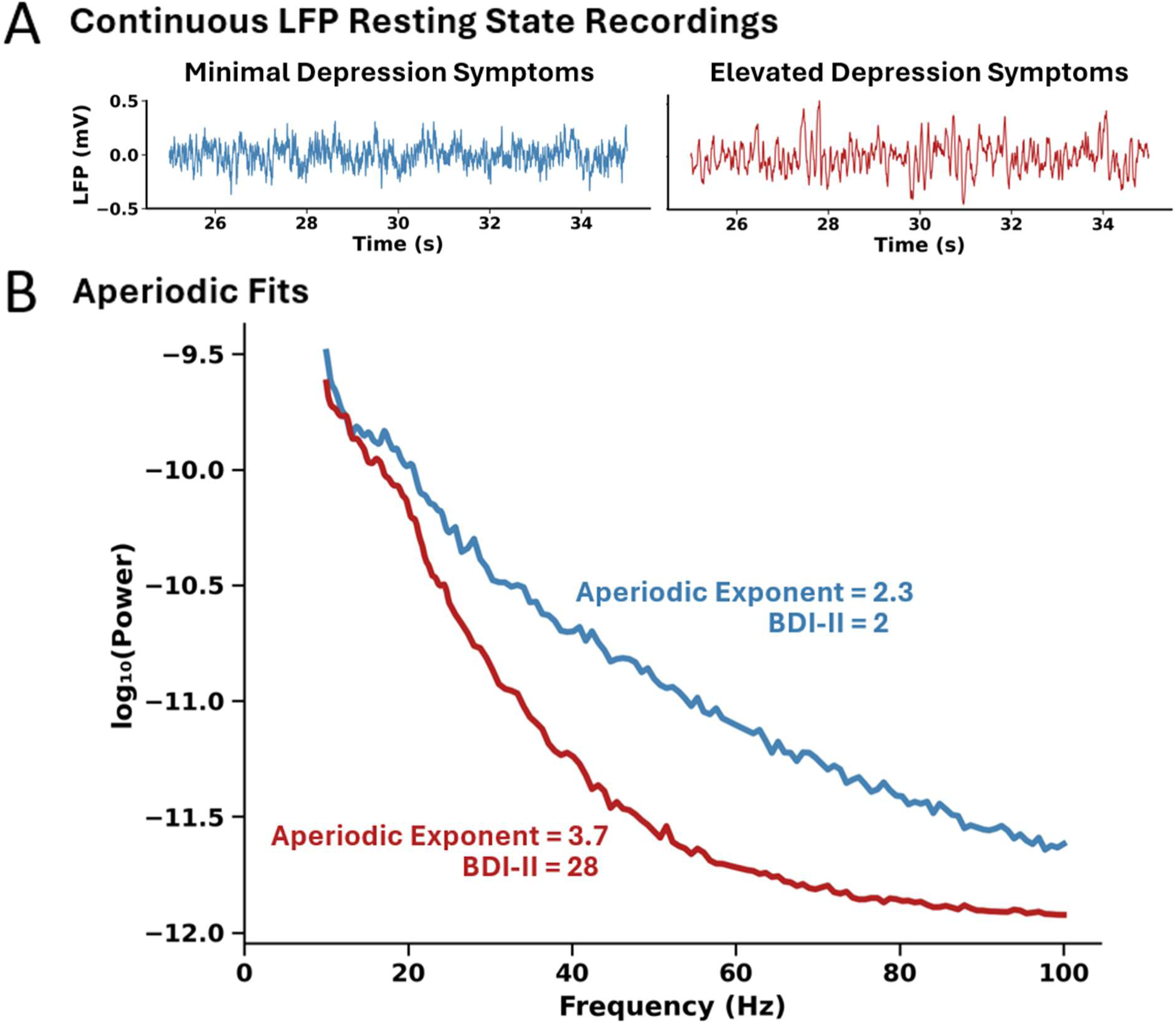
Derivation of the 1/f aperiodic exponent from intracranial LFP. A: Illustration from two representative left OFC LFP segments from one participant experiencing minimal depression symptoms (left) and one participant experiencing elevated depression symptoms (right). For each intracranial electrode, the power spectral density (PSD) was computed, and the aperiodic component was fit using the FOOOF algorithm^39^ to quantify the aperiodic exponent. Bottom: The PSD from the minimal symptoms participant (blue) has a flatter slope (smaller exponent) than that of the elevated symptoms participant (red), whose PSD exhibits steeper (larger exponent) activity.

### Group Comparisons of Aperiodic Exponent Values

To visualize group-level differences in aperiodic exponent values, we rendered electrode MNI coordinates in Visbrain^37^ separately for the minimal depression symptom and elevated depression symptom groups (**Fig. 3**). Additionally, whole-brain distributions were summarized with 30-bin count histograms and kernel-density overlays, and group-level differences were formally assessed using a two-sample Kolmogorov–Smirnov test with 20,000 permutations of group labels at the subject level.

### Discriminative Value of Aperiodic Exponent for Depression Symptom Severity Classification

To assess whether the aperiodic exponent differentiates participants experiencing minimal versus elevated depression symptoms (as defined by the BDI-II), we performed participant-level classification analyses at multiple spatial scales. First, we computed the whole-brain mean aperiodic exponent for each participant across all intracranial contacts, irrespective of anatomical region. Receiver operating characteristic (ROC) curves were constructed to quantify how well the whole-brain mean aperiodic exponent discriminated between participants with minimal versus elevated depression symptom severity, and the area under the curve (AUC) was calculated. The optimal decision threshold was defined using Youden’s J statistic. Classification performance was evaluated by the number of participants correctly and incorrectly classified (**Supp. Fig. 2**).

**Figure 2.**
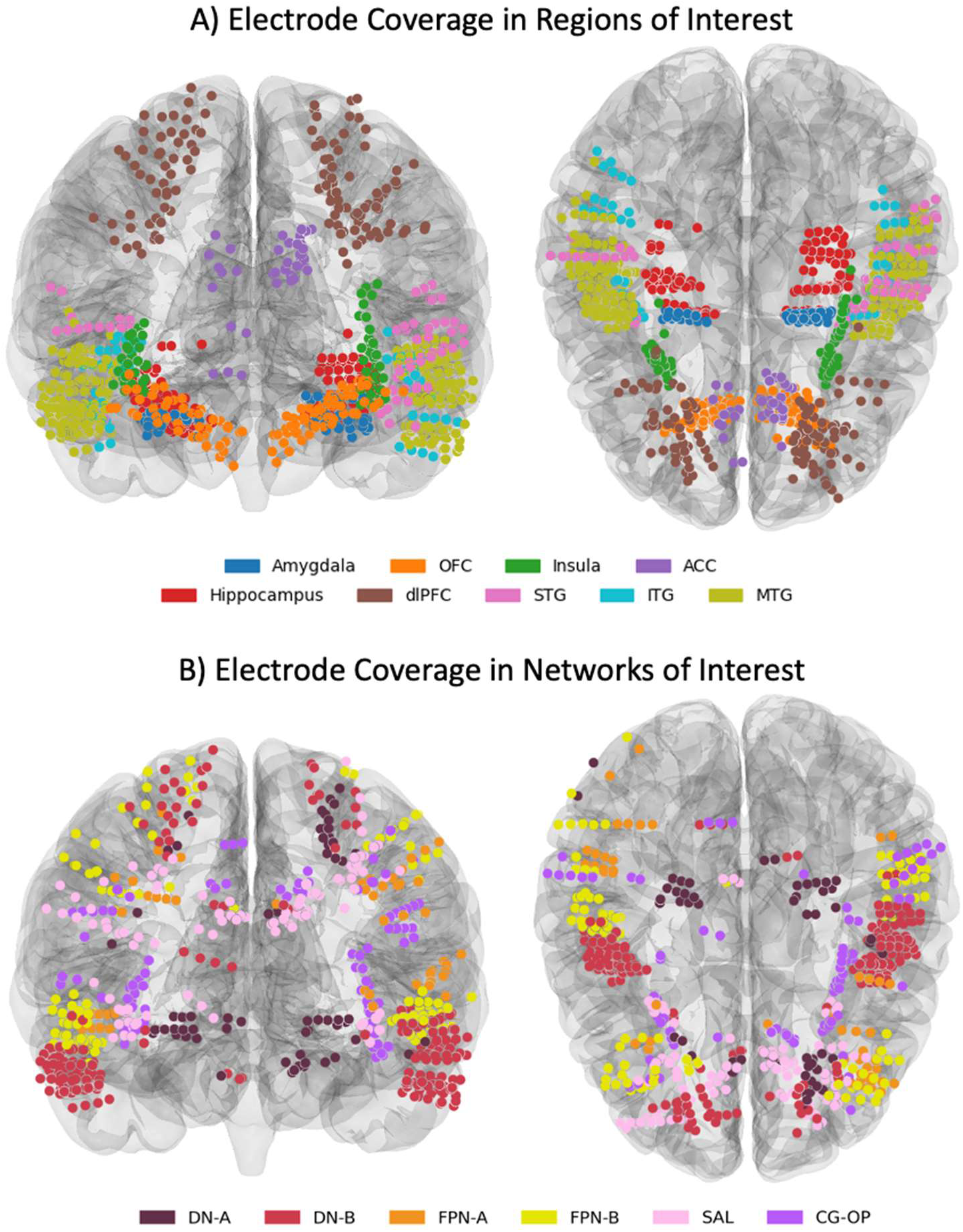
Intracranial electrode coverage. All participant electrode locations were mapped to a standardized cortical surface. A) Electrodes are color-coded by region: Orbitofrontal Cortex (OFC), Anterior Cingulate Cortex (ACC), Dorsolateral Prefrontal Cortex (dlPFC), Amygdala, Hippocampus, Insula, Superior Temporal Gyrus (STG), Middle Temporal Gyrus (MTG), Inferior Temporal Gyrus (ITG). B) Electrodes are color-coded by network of interest: Default Network-A (DN-A), Default Network-B (DN-B), Frontoparietal Control Network-A (FPN-A), Frontoparietal Control Network-B (FPN-B), Salience (SAL), Cingulo-Opercular (CG-OP). The distribution across participants highlights dense sampling within frontal, limbic, insular, and temporal regions, enabling region- and network-specific analyses of spectral aperiodic activity.

**Figure 3.**
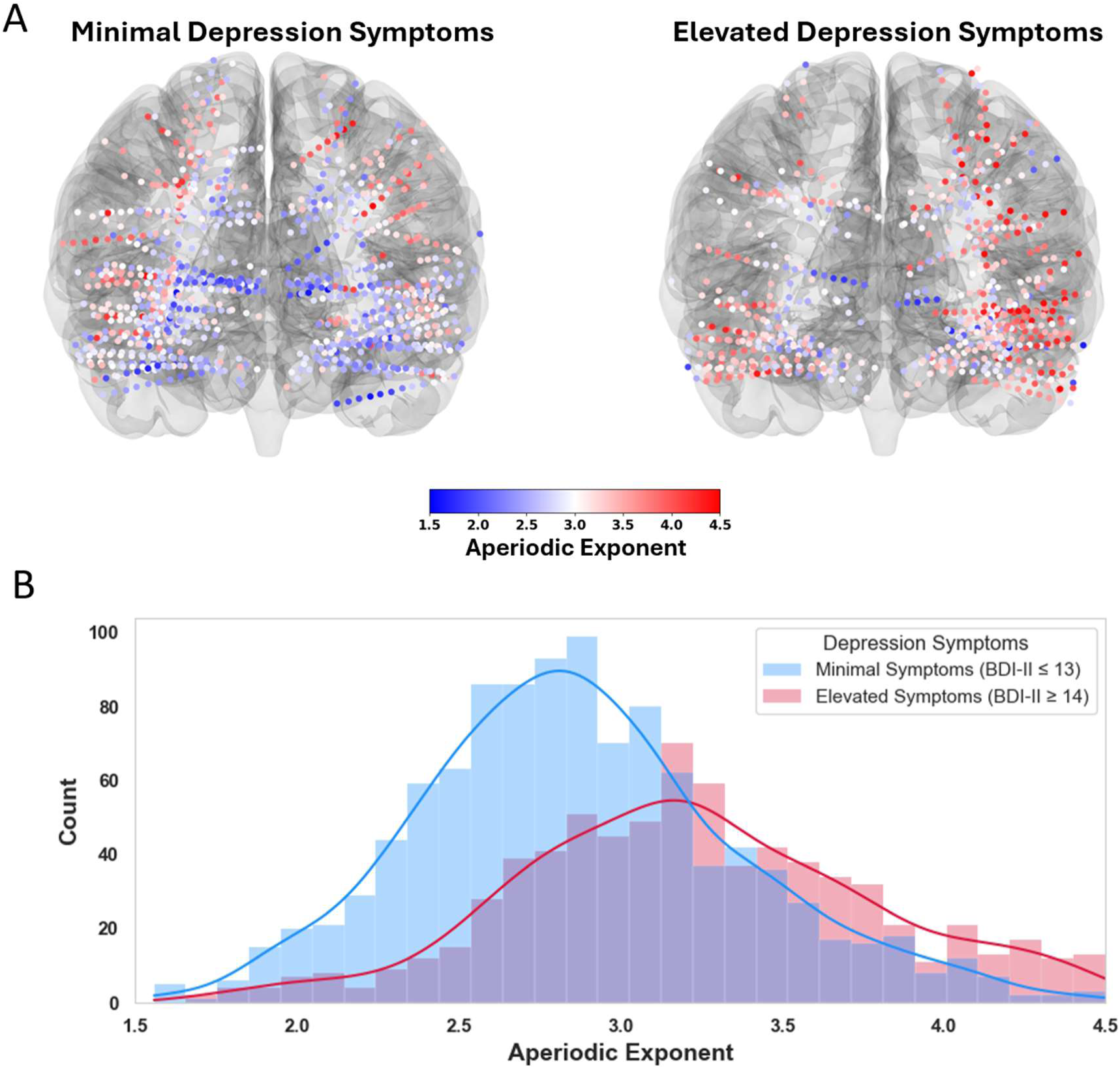
Global trends in aperiodic exponent separate minimal vs elevated depression symptom severity groups. A) Spatial maps of aperiodic exponent across all intracranial contacts, stratified by symptom burden: minimal depression symptoms (left) and elevated depression symptoms (right). Each dot marks a contact rendered on a template cortex; warmer colors (red) indicate higher exponent values and cooler colors (blue) indicate lower values (color scale 1.5-4.5). B) Histograms show aperiodic exponent value counts across all contacts for minimal (blue) and elevated (red) depression symptom groups with kernel-density overlays; y-axis indicates the number of contacts in each bin.

Second, identical ROC analyses were conducted separately for each region to identify region-specific contributions to discriminative performance. Regions were screened (for classification only) using Welch’s t-tests with a criterion of uncorrected p < 0.05. Finally, using only those regions that passed this screening, we evaluated whether an aggregated regional exponent value provided maximal discriminatory utility (post hoc; region screening and thresholding (Youden’s J) performed in-sample on the same dataset). Valid-region counts were compared between groups using a two-sided Mann–Whitney U test with rank-biserial effect size. Because screening and thresholding were performed on the same dataset, this aggregated regional analysis is an in-sample derivation rather than an out-of-sample validation. For this aggregated regional model, we report AUC, the selected cutoff, and bootstrap 95% confidence intervals (10,000 iterations) to characterize uncertainty.

### Calculating Region- and Network-Level Associations between Aperiodic Exponent and Depression Symptom Severity

To examine regional and network associations between the aperiodic exponent and depression symptom severity, electrode-level data were then aggregated to participant-level mean exponents for each region or network and modeled via ordinary least squares (OLS) regression against BDI-II total score, adjusting for age, given age’s relationship to the aperiodic component.^43^ Due to the sample size and gender distribution across symptom groups (minimal depressive severity: 4 F / 7 M; elevated depressive severity: 5 F / 4 M), gender was not included as an additional covariate in the primary analysis to avoid model overfitting and unstable estimates (See Supplementary Materials for full demographic analysis).

Model formula in Wilkinson notation:

(1) BDI-II Total Score ∼ region or network Exponent + Age

For each region, we also tested associations with BDI-II subscale scores (Somatic-Affective, Cognitive, and Anhedonia). At the network level, BDI-II subscale analyses were restricted to testing the relationship between SAL exponents and Anhedonia, given our *a priori* hypothesis that SAL would track Anhedonia.^21^

Model formula in Wilkinson notation:

(2) BDI-II Subscale Score ∼ region or network Exponent + Age

For each model, we extracted the exponent term’s standardized coefficient (βstd), and the partial correlation (partial r). Exponent-term p values were corrected for multiple comparisons across regions (and, separately, across networks) using the Benjamini–Hochberg FDR procedure; figures display best-fit regression lines per region or network with 95% CIs, with significant associations after FDR correction (pFDR) marked.

Additionally, we display region × subscale matrices of partial r with significance stars based on pFDR values.

### Leave-One-Subject-Out Robustness Analyses

To evaluate the stability of regional associations between aperiodic exponent and symptom measures, we performed a leave-one-subject-out (LOSO) robustness procedure for each region or network and outcome (BDI-II total and relevant subscales). For each model, we first fit an OLS regression predicting symptom score from region (or network)-mean exponent and age. The model was then iteratively refit after excluding one participant at a time, and the proportion of iterations that remained statistically significant (p < 0.05) was computed as the LOSO survival rate, quantifying robustness to participant removal.

Associations were operationally defined as robust if at least 80% of LOSO iterations remained significant. Results were summarized as a heatmap displaying LOSO survival percentages across regions (or networks) and outcomes (**Fig 6, 7)**.

## Results

### Participant coverage and signal characterization

We first visualized the anatomical distribution of intracranial electrodes across participants (**Fig. 2**). Electrodes were localized to nine cortical and subcortical regions (OFC, ACC, dlPFC, insula, amygdala, hippocampus, STG, ITG, and MTG) and six cortical association networks (DN-A, DN-B, FPN-A, FPN-B, SAL, CG-OP). Together, electrodes encompass key frontal, limbic, insular, and temporal zones thought to be implicated in depression.

### Aperiodic exponent differentiates participants by depression symptom severity

To test whether aperiodic exponent values differed between participants with minimal versus elevated depression symptom severity, we compared the whole-brain distribution of aperiodic exponent values across all recorded contacts (**Fig. 3**). Exponent values were projected onto a standardized cortical surface to visualize spatial differences between groups (**Fig. 3A**). A two-sample Kolmogorov-Smirnov test indicated a significant divergence in overall exponent value distributions between groups (D = 0.302, permutation p = 0.009; **Fig. 3B**).

To determine whether aperiodic exponent values could effectively discriminate depressive symptoms, we examined average exponent values at three levels per participant: (1) the whole-brain mean exponent (2) the mean exponent within each region, and (3) a combined regional model using regions screened as candidates by group differences in step 2 (using uncorrected p-values).

For the whole-brain analysis, an optimal cutoff of 3.00 was identified (t = 2.62, p = 0.020, AUC = 0.82), with three participants in the non-depressed group and two in the depressed group misclassified (**Supp. Fig 2**).

### Aperiodic exponents in OFC, ACC, insula, and amygdala show the strongest class separation

At the regional level, differentiation between groups was observed in the ACC (p = 0.004, t = 3.47, d = 1.68), OFC (p = 0.029, t = 2.50, d = 1.18), insula (p = 0.032, t = 2.40, d = 1.24), and amygdala (p = 0.005, t = 3.52, d = 1.71) (**Table 2**). All p values are uncorrected for multiple comparisons and were used only for regional screening. These region-level results are post-hoc and should be interpreted as exploratory.

**Table 2.** Region-wise discrimination and group differences in aperiodic exponent. For each region (left column), AUC quantifies how well the aperiodic exponent separates participants experiencing minimal depression symptom severity (MS) from those with elevated depression symptom severity (ES) groups. Cutoff is the decision threshold used for classification. The “p, t” column reports the two-tailed Welch’s t statistics (t) and uncorrected p value (p) for the group difference in exponents. Cohen’s d gives the standard effect size derived from the t and the group sample sizes. “ES, MS” lists the number of unique participants contributing data to each group. “Miscl. ES” and “Miscl. MS” are the numbers of misclassified minimal and elevated depression symptom participants at the listed cutoff. Cell shading: in “p, t,” green indicates p < 0.05; in Cohen’s d, green ≥ 1.15, yellow 0.60–1.14, red < 0.60; in Miscl. columns, green = 0 errors, yellow = 1–2, red ≥ 3.

| Region | AUC | Cutoff | p, t | Cohen's d | ES, MS | Misc. ES | Misc. MS |
| --- | --- | --- | --- | --- | --- | --- | --- |
| ACC | 0.87 | 2.82 | $p = 0.004$, $t = 3.47$ | 1.68 | 7, 10 | 0 | 2 |
| STG | 0.69 | 2.88 | $p = 0.346$, $t = 1.01$ | 0.58 | 6, 6 | 0 | 3 |
| OFC | 0.81 | 2.91 | $p = 0.029$, $t = 2.50$ | 1.18 | 8, 10 | 1 | 1 |
| Hippocampus | 0.77 | 2.77 | $p = 0.079$, $t = 1.88$ | 0.89 | 7, 11 | 1 | 4 |
| ITG | 0.83 | 3.10 | $p = 0.123$, $t = 1.76$ | 1.02 | 6, 6 | 1 | 0 |
| Insula | 0.84 | 2.85 | $p = 0.032$, $t = 2.40$ | 1.24 | 8, 7 | 2 | 1 |
| MTG | 0.75 | 3.27 | $p = 0.192$, $t = 1.38$ | 0.62 | 9, 11 | 2 | 2 |
| Amygdala | 0.86 | 2.80 | $p = 0.005$, $t = 3.52$ | 1.71 | 8, 9 | 3 | 0 |
| dLPFC | 0.63 | 3.43 | $p = 0.076$, $t = 1.91$ | 0.85 | 9, 11 | 5 | 1 |

Finally, in a post-hoc, in-sample analysis, we combined the average exponent values from the ACC, OFC, insula, and amygdala. The number of valid regions per subject did not differ significantly between depression groups (Mann–Whitney U = 43.5, p = 0.647, rank-biserial r = 0.121). This combined measure yielded a cutoff of 2.91 (AUC = 0.86, 95% CI = 0.64-1.00) producing excellent in-sample separation with one participant from each group being misclassified (**Fig. 4**).

**Figure 4.**
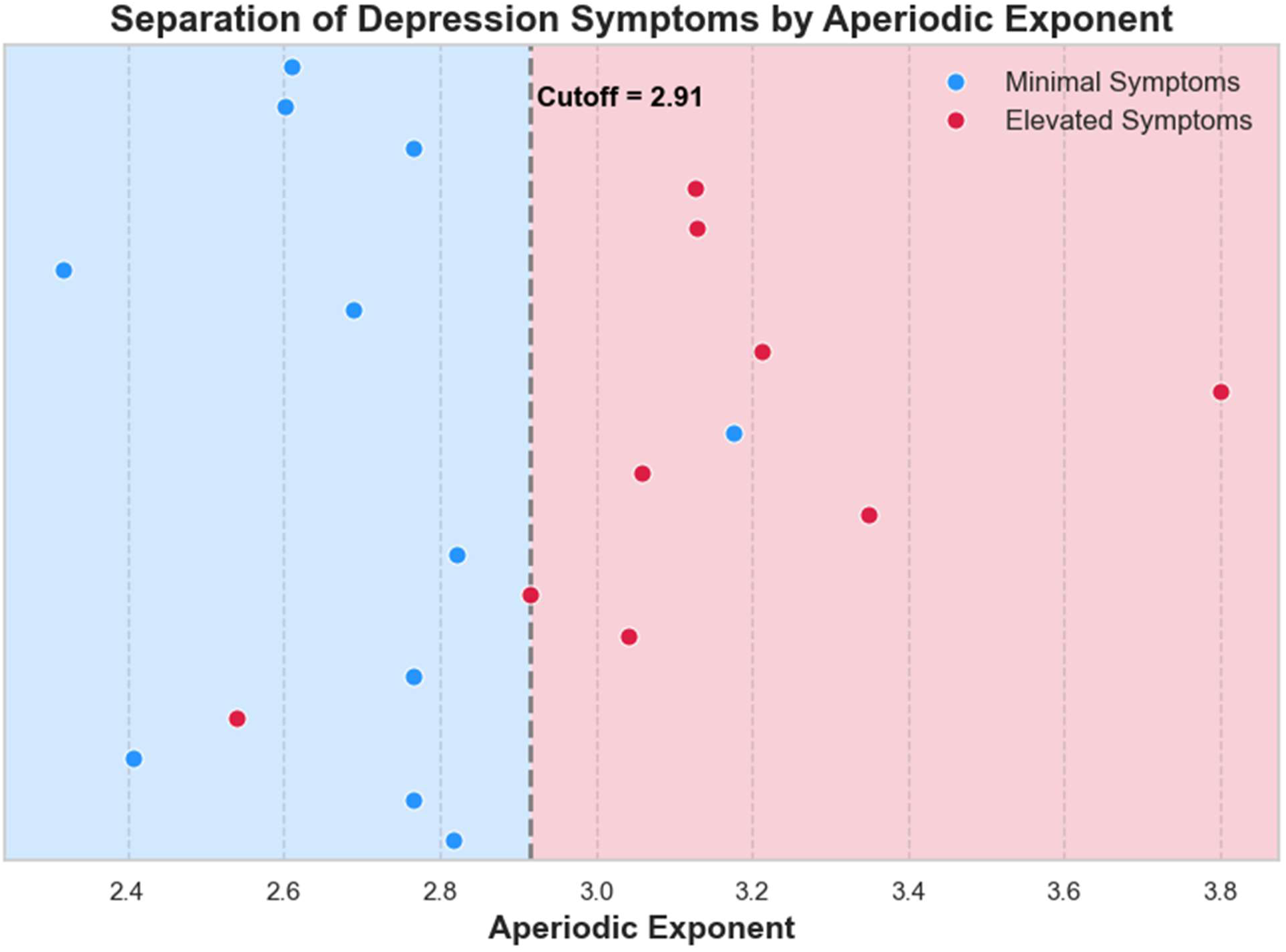
Participant-level separation of symptom status by aggregated OFC, ACC, insula, and amygdala aperiodic exponent values. Each dot is one participant (mean exponent across the four regions). Points are colored by true symptom status (blue = minimal depression symptoms; red = elevated depression symptoms). The dashed vertical line marks the optimal Youden’s J threshold (cutoff = 2.91); blue/red shading indicates the class assignment under this threshold. Note that region-wise discrimination and group-difference analyses are post-hoc and in-sample; cutoffs and CIs were derived on the same dataset and are hypothesis-generating, not validation. * = outlier in 80–95 Hz high-gamma coherence (putative artifact; see **Supp.** Fig 1).

### Aperiodic exponent tracks BDI-II (total & subscales) in specific regions and networks

We next examined whether aperiodic exponent values within each region or network scaled continuously with depressive severity across participants. To assess this relationship, we fit OLS regression models for each region and each network, relating mean exponent values to total BDI-II scores, while controlling for age (**Fig. 5**). In line with our post-hoc classification analysis (**Fig. 4**), the same four regions showed a significant positive correlation between BDI-II scores and mean exponent values after correcting for multiple comparisons: ACC (pFDR = 0.019, βstd = 0.63, partial r = 0.65), OFC (pFDR = 0.019, βstd = 0.66, partial r = 0.69), insula (pFDR = 0.019, βstd= 0.68, partial r = 0.70), and amygdala (pFDR = 0.027, βstd = 0.68, partial r =0.61). For the cortical association networks, SAL (pFDR = 0.024, βstd = 0.62, partial r = 0.63) and DN-B (pFDR = 0.046, βstd = 0.56, partial r = 0.55) showed significant positive correlations between BDI-II scores and network-averaged exponent values.

**Figure 5.**
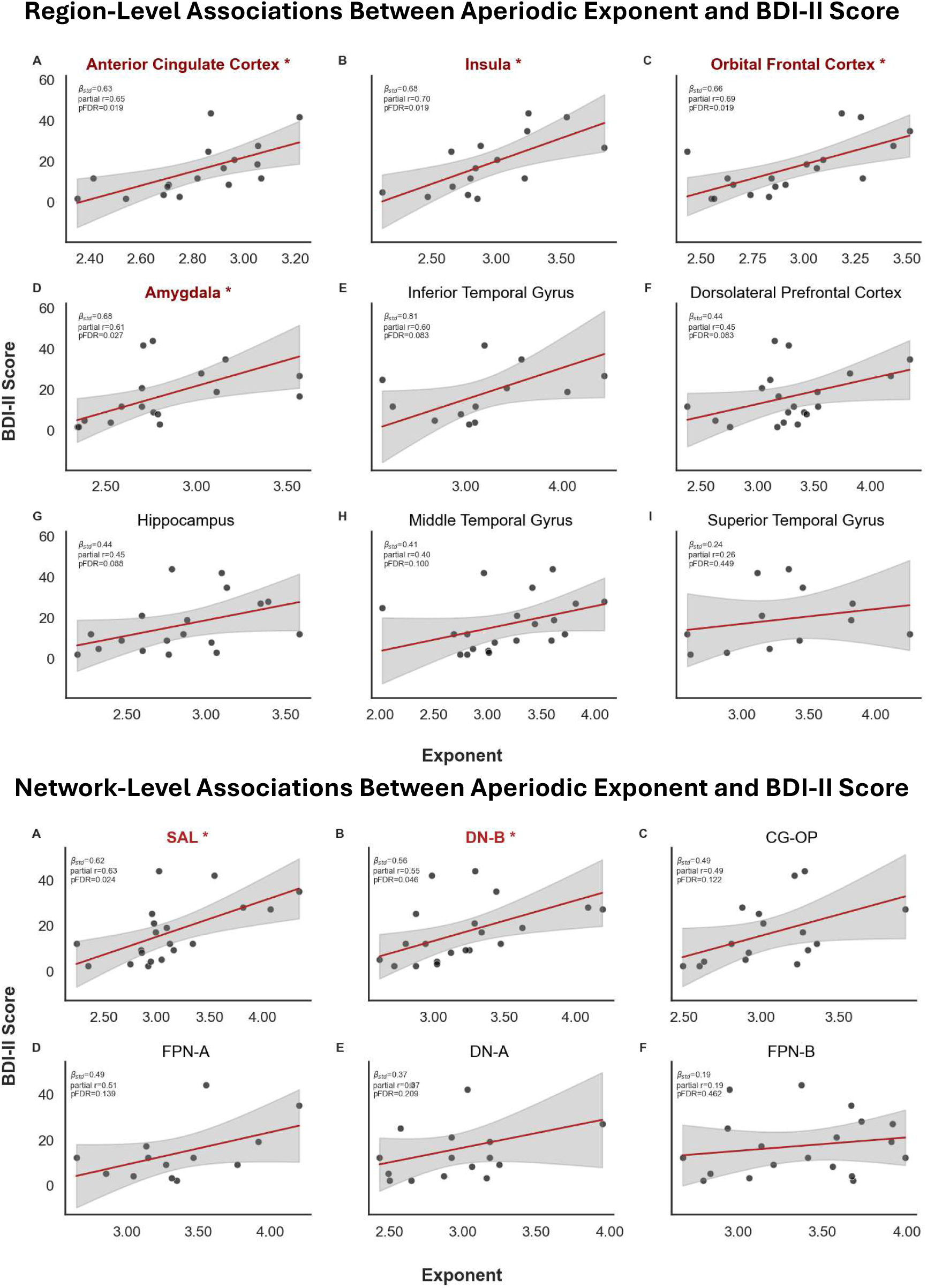
Regional and network level associations between aperiodic exponent and continuous BDI-II score. Top Panel: Regional comparisons. Bottom Panel: Network comparison. Each subplot shows one region/network; each point is a participant level mean (contact values within that region/network collapsed to one value per participant). The red line is the OLS fit of BDI-II score on exponent adjusting for Age (BDI-II Score ∼ Exponent + Age), with the shaded 95% CI. Insets report the partial r, standardized coefficient (βstd), and the Benjamini–Hochberg FDR–adjusted p-value (pFDR) for that panel. Axes: x = aperiodic exponent; y = BDI-II score.

**Figure 6.**
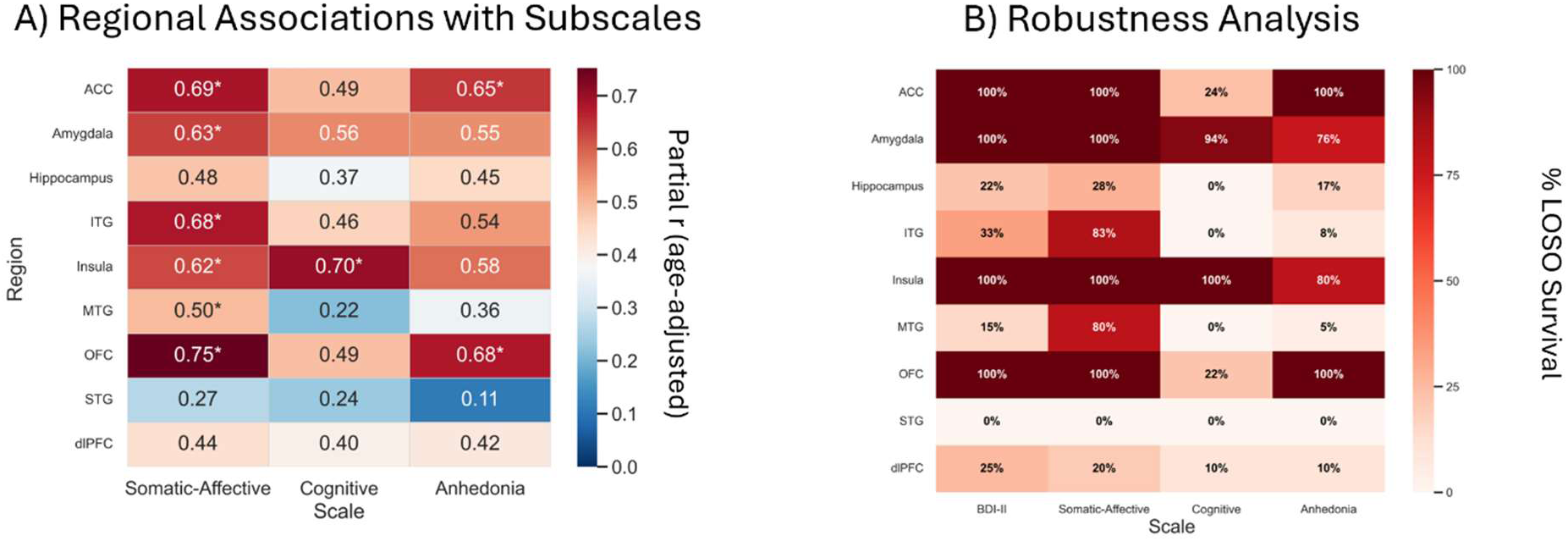
Regional aperiodic exponent associations with depression symptoms: Subscale effects and participant-level robustness. A) Heatmap of partial r between regional aperiodic exponent and BDI-II subscales; asterisks mark FDR-corrected p < 0.05, color encodes the magnitude of partial r. B) Heatmap showing the percentage of LOSO refits in which the aperiodic exponent retained a significant association with total BDI-II score and each BDI-II subscale within each region (α = 0.05). Cell annotations report percent survival; color spans 0-100%, with higher values indicating more robust effects across participants).

### Regional associations with subtypes of depression symptoms

To explore potential regional specificity in how the aperiodic exponent relates to subtypes of depressive symptomatology, we examined the association between region-mean exponent values and BDI-II subscale scores (Cognitive, Somatic-Affective, and Anhedonia), using the same OLS regression framework, while controlling for age (**Fig. 6A**). Given that the subscales were derived from the same BDI-II scale used for the previous analyses, these analyses were considered exploratory.

Significant positive associations between subscale and aperiodic exponent values were observed between Somatic-Affective symptoms and the OFC (pFDR = 0.004), ACC (pFDR = 0.015), amygdala (pFDR = 0.026), insula (pFDR = 0.037), ITG (pFDR = 0.037), and MTG (pFDR = 0.047). For cognitive symptoms, the insula showed a significant positive association (pFDR = 0.044). For anhedonia, the OFC (pFDR = 0.025) and ACC (pFDR = 0.031) showed significant positive associations.

### Regional exponent–symptom associations remain robust under leave-one-subject-out validation

To evaluate the stability of regional associations between aperiodic exponent values and depressive symptom measures, we performed a LOSO robustness analysis for each region and outcome (BDI-II total score and its Cognitive, Somatic-Affective, and Anhedonia subscales; **Fig. 6B**). For each OLS model, we computed the proportion of iterations where the exponent-symptom relationship maintained significance (p < 0.05) after exclusion of one participant.

Consistent with the primary OLS regression analyses, the ACC, OFC, insula, and amygdala exhibited robust positive associations with total BDI-II scores, with 100% survival in these four regions. The Somatic-Affective subscale showed a similar pattern of robustness, with 100% survival in the ACC, OFC, insula, amygdala, 83% in the ITG, and 80% in the MTG. The Cognitive subscale showed robust effects in the insula (100%) and amygdala (94%), while the Anhedonia subscale showed robustness in the OFC (100%), ACC (88%), and insula (80%).

Together, these results indicate that the primary region exponent-symptom associations—particularly in fronto-limbic regions—are not driven by individual participants and remain consistent across subsamples.

### A priori SAL–anhedonia association and BDI-II network effects withstand LOSO

Previous literature suggests that SAL predicts future symptoms of anhedonia and likely overlaps with modern neuromodulation targets.^21–23^ As such, we predicted that SAL aperiodic exponents would be greater for participants with increased Anhedonia symptoms. Critically, we observed a significant positive association between anhedonia symptom scores and SAL aperiodic exponents (**Fig. 7A**; p = 0.004, βstd=0.63, partial r = 0.62). To evaluate the stability of network-level associations between aperiodic exponent values and depressive symptom measures, we performed a LOSO robustness analysis for each network and the BDI-II total score (**Fig. 7B**). Network effects were robust: SAL and DN-B vs total BDI-II score and the SAL–Anhedonia subscale association each exhibited 100% LOSO survival.

**Figure 7.**
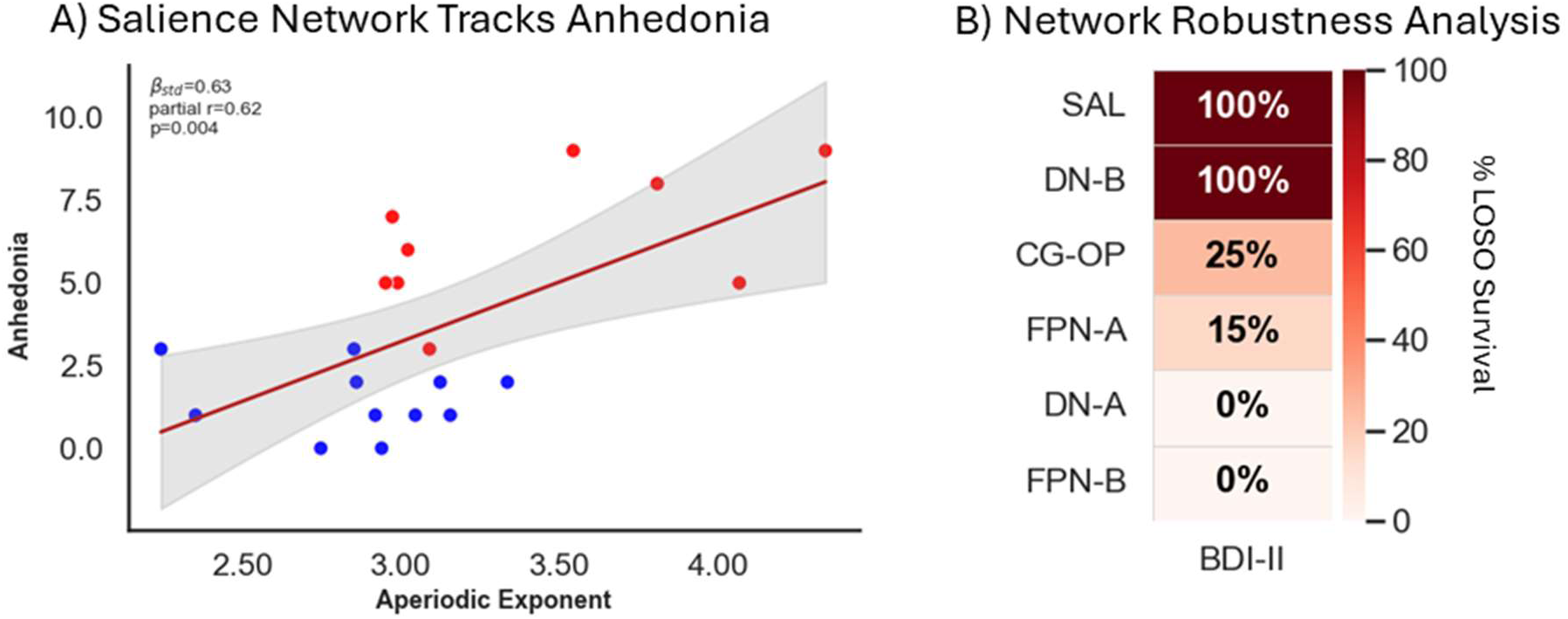
Network aperiodic exponent associations with depression symptom severity: Salience-Anhedonia and regression participant-level robustness (LOSO). A) Relationship between the aperiodic exponent in the Salience Network and the Anhedonia subscale. Each point is one participant’s aperiodic exponent mean value within SAL (color coded to mood status: blue = minimal depression symptoms, red = elevated depression symptoms). The line is the OLS fit of Anhedonia score on exponent adjusting for Age (Anhedonia Score ∼ Exponent + Age), with the shaded 95% CI. B) Percentage of LOSO refits in which the aperiodic exponent remained significantly related to total BDI-II score (α = 0.05). Cell annotations report percent survival: color spans 0-100%, with higher values indicating more robust effects across participants.

## Discussion

MDD lacks objective neural markers, constraining efforts to monitor treatment progress as behavioral measures of depression often emerge after weeks or months of therapy. To address this gap, we leveraged intracranial resting-state recordings from patients with drug-resistant epilepsy, which provided rare access to frontal, limbic, and insular structures implicated in depression.

In this study, we focused on the non-oscillatory component of the neural power spectra to compute the aperiodic exponent (spectral slope)—a compact and passive metric thought to index aspects of broadband excitability and, by extension, excitation-inhibition balances.^14, 15^ Emerging work indicates that alterations in this background spectral trend carry physiological relevance in psychiatric illness, including MDD.^13^ Accordingly, we quantified exponents in the 10-100 Hz range, which yields stable aperiodic estimates in iEEG^40^ and encompasses the canonical pattern observed with reductions in depressive symptoms: decreased low-frequency and increased high-frequency power. This pattern is reflected in a flattening of the spectral slope (lower aperiodic exponent).

To localize these spectral changes, we focused on regions with adequate sampling across participants, including the OFC, dlPFC, ACC, amygdala, hippocampus, insula, ITG, STG, and MTG. Additionally, we investigated cortical association networks thought to be implicated in MDD, including the salience network and default network. This approach allowed us to assess how the aperiodic component of neural power spectra, in distinct anatomical regions and networks, underlies variation in depression symptom severity.

Consistent with our hypothesis, higher aperiodic exponent values, representing greater low- relative to high-frequency activity, were associated with greater depression symptom severity.

### Higher whole-brain aperiodic exponents track depressed mood

We found that the whole-brain distribution of aperiodic exponent values distinguished participants with minimal depression symptoms (BDI-II ≤ 13) from those with elevated depression symptoms (BDI-II ≥ 14). These global patterns align with large-scale neuroimaging evidence that characterizes MDD by inefficient cortical processing and disrupted network integration: a consortia-scale dataset on MDD demonstrated widespread reductions in functional connectivity relative to healthy controls.^44^ Within this framework, a higher aperiodic exponent could reflect reduced broadband excitability and serve as a candidate electrophysiological correlate of inefficient cortical processing in MDD.

### Fronto-limbic and insular aperiodic exponents preferentially track depressed mood

While whole-brain mean exponent values discriminated participants experiencing minimal versus elevated depression symptoms, region-specific effects were even more pronounced. Exponents scaled with BDI-II scores only in the OFC, ACC, insula, and amygdala, indicating that, within this cohort, the symptom-tracking signal is preferentially expressed in these four regions. A post-hoc multiregional composite spanning these four regions captured most of the discriminative information and yielded the strongest group separation. Interestingly, these regions appear to participate in the functional circuits that underlie currently utilized deep brain stimulation targets for treatment-resistant depression.^8^

Across BDI-II depression subscales, our exploratory findings suggest that somatic–affective burden may map to fronto-limbic/temporal zones,^45, 46^ cognitive features implicate the insula,^47, 48^ and anhedonia trends align with ACC/OFC reward circuits;^21, 49^ these patterns are consistent with prior work, and are presented here strictly for hypothesis-generating purposes.

These associations, for both total BDI-II and the subscale scores, were robust under LOSO analyses, indicating that no single participant drove the effects.

### Salience and Default network aperiodic exponents preferentially track depression symptom severity

When applying a network-based approach to the aperiodic exponent, we found that two cortical association networks, the salience (SAL) and default (DN-B) networks, were significantly associated with depression symptom severity. This pattern aligns with the regional associations identified above, as SAL encompasses regions of the ACC and anterior insula^50^ and DN-B is coupled to the amygdala.^51^ Default-mode circuitry has been linked to self-referential processing and rumination in MDD.^52^ SAL supports reward processing and detection of novel, behaviorally relevant stimuli.^53, 54^ Our findings also converge with computational models suggesting SAL is a relevant target in clinical neuromodulation protocols for depression.^22^ Taken together, the increased aperiodic exponent we observe within these networks suggests that depression involves not only altered large-scale network organization but also a systematic change in cortical excitability.

Notably, we also found that aperiodic exponents in SAL significantly correlated with anhedonia symptoms, consistent with prior work showing that functional connectivity within SAL predicts future anhedonia symptoms.^21^ Prior meta-analyses on depression have also reported aberrant functional correlations between regions in the salience network and default network.^55^ Our observations on the aperiodic exponent in SAL and DN-B, robust to LOSO analyses, provide an electrophysiological validation to the fMRI-based functional connectivity literature.

### Towards a brain-based biomarker for depression symptom severity

Although whole-brain distributions of aperiodic exponent values distinguished participants experiencing minimal versus elevated depression symptoms, the most informative signal was regionally specific. This pattern aligns with evidence that whole-brain connectivity metrics are modest biomarkers of depression, whereas more spatially specific circuit features carry greater discriminative value.^56, 57^ Exponents scaled with BDI-II scores in the OFC, ACC, insula, and amygdala, and a post hoc composite of these four regions captured most of the discriminative information and yielded the strongest group separation. In a convergent network-level analysis, SAL and DN-B also scaled with BDI-II scores, two systems that help assign value to external and internal stimuli and shape other-directed thought, respectively. Therefore, our findings suggest that depression symptoms are accompanied by an increase in the aperiodic exponent within fronto-limbic and insular regions consistent with a shift in local cortical excitability toward a more low-frequency-dominated (higher exponent) state. In this framework, the aperiodic exponent provides a scalable, physiologically interpretable readout of low- versus high-frequency power balance and could be used as a continuous brain biomarker of treatment target engagement and circuit-level response, supporting future pharmacological and neuromodulation interventions that aim to normalize aberrant dynamics in these circuits and ultimately improve mood. Thus, the aperiodic exponent, in tandem with behavioral measures, offers a physiological component of symptomatology that may track treatment effects on a timescale more closely aligned with intervention decisions. As such, we view this work as an initial step toward clinically actionable brain biomarkers of depression symptom severity, and future mood decoding.

## Limitations and future directions

While the present results are robust across analyses (including LOSO robustness checks), limitations qualify their interpretation. First, the cross-sectional design precludes causal inference. We partially mitigated this by administering the BDI-II immediately before the resting-state recording to obtain a temporally relevant readout of depression symptoms. Yet the biomarker-symptom time constant, and any actionable lead-lag for intervention in neuromodulation, remains unknown. Secondly, we focused on participants’ present depression symptom status. Accordingly, we did not incorporate lifetime clinical history of depression or diagnostic information into the analyses. We view this emphasis on current symptom burden as a strength allowing for broader applicability because it is not dependent on a specific clinical diagnosis. Longitudinal and interventional studies are needed to test whether exponent changes precede depression symptoms and to determine when to intervene (onset vs predictive states) and how to titrate (dose, duration, and frequency). Because regional screening and thresholding were performed on the same data used for evaluation, the combined regional AUC is an apparent in-sample estimate and likely optimistic; external validation is needed. Additionally, participants were patients with epilepsy. Although seizure-onset-zone contacts were excluded, cohort factors such as medication burden may limit generalizability; nevertheless, the high prevalence of comorbid depression symptoms in epilepsy lends ecological relevance to the sample.

Lastly, spatial specificity can be influenced by region and parcellation choices; replication across parcellations and cohorts will clarify generalizability. Still, the biological meaning of increased aperiodic exponents requires causal tests; longitudinal and interventional work (e.g., TMS/DBS/ECT/ketamine) will be essential to determine whether a multiregional or network exponent composite can complement symptom scales for treatment monitoring. These caveats outline future directions rather than detract from the core finding that aperiodic exponents in fronto-limbic and insular circuits provide a sensitive readout of depression symptom severity.

## Conclusion

Across the OFC, ACC, insula, and amygdala, and at the level of the SAL and DN-B cortical association networks, intracranial aperiodic exponents reliably differentiated and tracked depression symptom severity, consistent with current systems-level views of MDD. By distilling low-to-high frequency power balance into a single, interpretable value, the exponent is well-suited for physiology-guided tracking of treatment response as behavioral symptom ratings accrue over weeks to months. In this cohort, effects were robust to LOSO analyses, suggesting stability at the participant level. Prior iEEG studies of the aperiodic exponent have largely been limited to treatment-resistant depression; here we show that the exponent–depression symptom severity relationship may generalize more broadly, supporting the exponent as a foundation for decoding depression status, and potentially other mood disorders. Looking ahead, integrating exponent tracking into therapeutic workflows (e.g., target selection, dose titration, closed-loop triggers) and testing on independent cohorts and non-invasive recordings will be essential to establish generalizability and to test whether exponent-based monitoring can meaningfully guide neuromodulation protocols.

## Data Availability

The data that support the findings of this study are available on request from the corresponding author.

